# Perivascular fibroblasts locally regulate the vasomotor dynamics of pial arterioles

**DOI:** 10.64898/2026.09.11.751046

**Authors:** Stephanie Karas, Elisabeth Harmon, Orla Bonnar, Bradley Pawlikowski, Danielle A. Jeffrey, Laura M. Muñoz, Fiona Lake, Maria J. Sosa Ibarra, Cara D. Nielson, Fabrice Dabertrand, Julie A. Siegenthaler, Andy Y. Shih, Stephanie K. Bonney

**Affiliations:** Department of Anesthesiology, University of Colorado Anschutz Medical Campus, Aurora Colorado, USA; UK Dementia Research Institute, University College London, London, United Kingdom; Department of Pediatrics, Section of Developmental Biology, University of Colorado Anschutz Medical Campus, Aurora Colorado, USA; Center for Developmental Biology and Regenerative Medicine, Seattle Children’s Research Institute, Seattle Washington, USA; Department of Pharmacology, University of Colorado Anschutz Medical Campus, Aurora Colorado, USA; Department of Pediatrics, University of Washington, Seattle Washington, USA

## Abstract

Perivascular fibroblasts (PVFs) ensheath pial arterioles on the brain surface, yet their physiological role on the healthy cerebrovasculature remains unknown. Here, we show that PVFs display calcium dynamics that are temporally aligned with arteriolar vasomotion. Depletion of fibroblasts disrupted vasomotion whereas enhancement of PVF-Gq signaling increased vasomotor activity and accelerated fluid influx into the peri-arteriolar space. These findings identify PVFs as unrecognized regulators of cerebral vasodynamics and cerebrospinal fluid movement.

## Main

Perivascular fibroblasts (PVFs) surround pial and penetrating arterioles of the brain, lining the outer layer of vascular smooth muscle cells (SMCs)^1^. They reside within the perivascular space with macrophages which acts as conduits for cerebrospinal fluid (CSF) influx and interstitial solute clearance^1–6^. Brain arterioles exhibit rhythmic diameter oscillations during rest, termed vasomotion^7^, which may drive perivascular fluid transport. However, whether PVFs contribute to arteriolar vasodynamics and fluid movement remains unknown.

To examine this possibility, we generated Col1a2CreER-iDTR mice and applied diphtheria-toxin (DTX) and AAV8-CAG-GFP to the exposed cortical surface during cranial window implantation to deplete and label brain fibroblasts^8–10^ (**Extended Data Fig. 1a, b**). Col1a2CreER recombines dural, leptomeningeal, and perivascular fibroblasts^4^, this approach depletes fibroblasts on the brain surface. By 7-days post DTX, awake Col1a2CreER-iDTR mice exhibited a marked reduction in GFP+ leptomeningeal fibroblasts (LPMFs) and PVFs, accompanied by disruption of arteriolar vasomotor dynamics, whereas controls retained fibroblasts and normal vasomotor rhythms (**Extended Data Fig. 1c-g**). ɑ-smooth muscle actin expression was preserved, indicating SMCs were maintained despite the disruption to vasomotion (**Extended Data Fig. 1h, i**).

We next asked whether PVFs respond to arteriole diameter changes. Since calcium (Ca^2+^) signaling in SMCs is coupled to arteriole diameter^11^, we tested whether PVFs also exhibit Ca^2+^ responses to vasomotricity. We isolated and pressurized parenchymal arterioles from Col1a2CreER-GCaMP6f mice and examined PVF-Ca^2+^ responses *ex vivo* following increased intraluminal pressure and K^+^-induced vasoconstriction^12,13^. Previously, we found that the Col1a2CreER mouse line, ∼4% of the recombined perivascular cells were SMCs and are readily distinguished from PVFs by their characteristic banded morphology compared to the flattened somata and thin, fibrillar processes of PVFs^4^ (**Extended Data Fig. 2**). Thus, this model enabled simultaneous measurement of Ca^2+^ activity in PVFs and sparse, morphologically distinct SMCs. Increasing intraluminal pressure in 20mmHg step increments and 60mM K^+^-induced vasoconstriction elicited rapid increases in both PVF- and SMC-Ca^2+^ which corresponded with vasoconstriction (**Fig. 1a-d**). This supports the physiological engagement of PVF-Ca^2+^ during vasomotricity.

**Figure 1.**
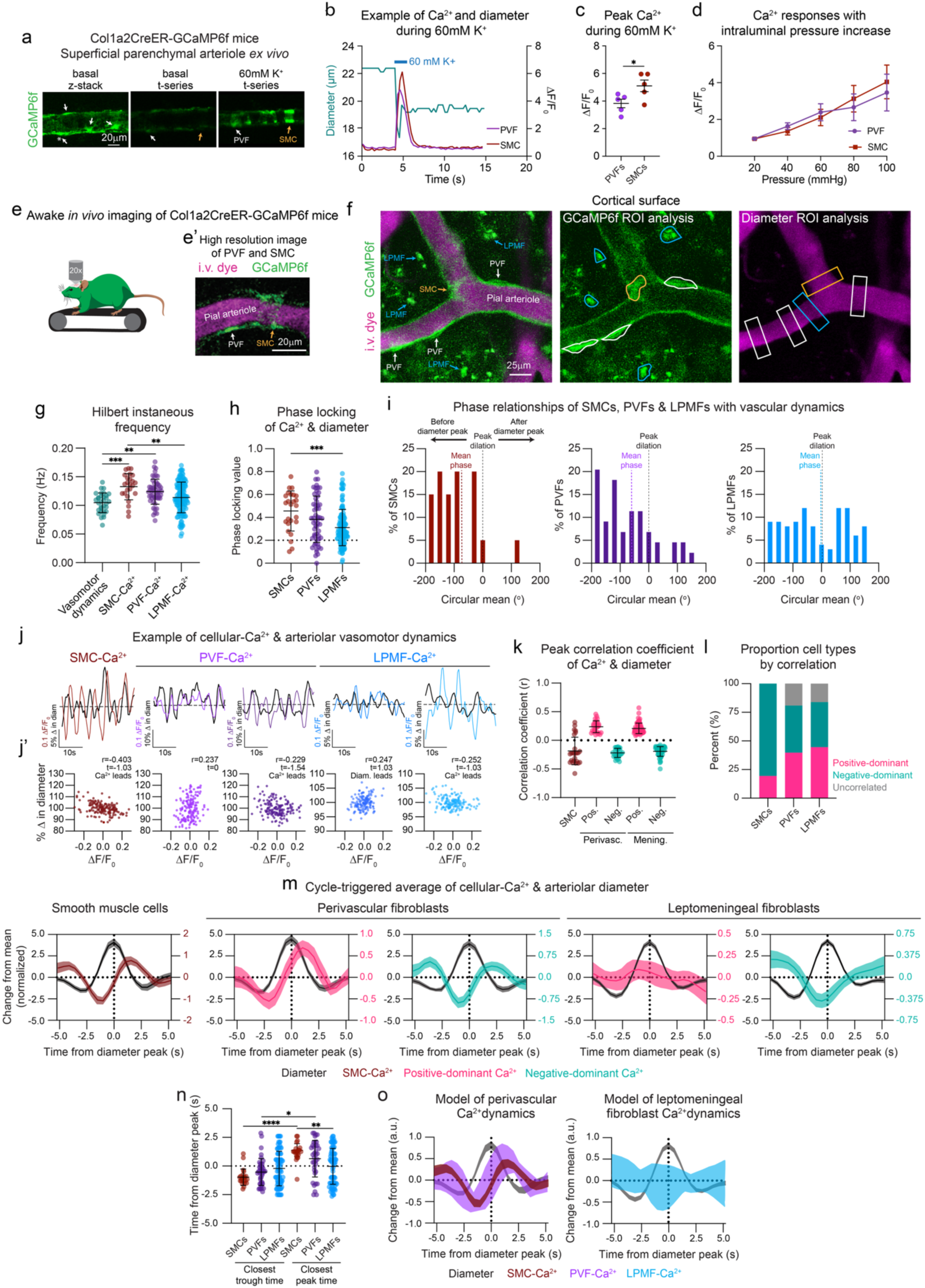
Perivascular fibroblast-Ca^2+^ activity is temporally coupled with resting arteriolar dynamics. **(a)** Representative *ex vivo* confocal images of a superficial parenchymal arteriole from Col1a2CreER-GCaMP6f mice with (left) z-stack, max projection taken under basal conditions, (middle) single-plane t-series under basal condition, and (right) single-plane t-series immediately following 60mM K^+^ exposure to the vasculature. GCaMP6f+ perivascular fibroblasts (PVFs; white arrows) are visible under basal conditions, with banding of recombined GCaMP6f-expressing smooth muscle cells (SMCs; blue arrows) visible after 60mM K^+^. Asterisk (*) indicates PVF that remained in the single-plane t-series for Ca^2+^ analyses. **(b)** Representative trace of a PA with 60mM K^+^ applied to the vasculature (blue line). Both diameter (μm, green) and change in Ca^2+^ (ΔF/F_0_) displayed for PVFs (purple, n=3) and SMCs (red, n=3). **(c)** Summary data of 60mM K^+^-induced increase in Ca^2+^ averaged per mouse (n=4 mice). Unpaired t-test, *p=0.0408. **(d)** Summary data of Ca^2+^ dynamics in response to increases in intraluminal pressure (20 to 100 mmHg) of PVFs and SMCs. Data were not significant per two-way repeated-measures ANOVA followed by Bonferroni’s correction post-hoc, n=4 mice. **(e)** Graphical depiction of awake *in vivo* imaging setup of Col1a2CreER-GCaMP6f mice head-fixed on freely-moving treadmill along with (**e’**) a representative high resolution two-photon images of GCaMP6f expression in PVFs (white arrow) and occasional SMCs (orange arrow) using the Col1a2CreER driver. PVFs and SMCs on pial arterioles can be distinguished from each other due to the banded, wrapping morphology of SMCs and flattened soma of PVFs aligned along the vascular wall. I.v. dye (70kDa Texas red-Dextran) shown in magenta with GCaMP6f in green. **(f)** Representative single-plane t-series *in vivo* image of a pial arteriole (magenta) on the cortical surface surrounded by GCaMP6f-expressing (green) PVFs (white), leptomeningeal fibroblasts (LPMF; blue), and SMCs (orange). Regions of interest (ROI) for measuring fluorescent intensity highlighted for each cell type in the 5-minute movies are shown in the GCaMP6f image. ROIs for measuring diameter changes in pial arterioles for the respective cell types are shown in the i.v. dye image. **(g)** Graphs of Hilbert instantaneous frequency (Hz) of arteriolar vasodynamics (green), SMC-Ca^2+^ oscillations (red), PVF-Ca^2+^ oscillations (purple), and LPMF-Ca^2+^ oscillations (blue). One-way ANOVA followed by Tukey’s multiple comparison test: Vasomotor dynamics vs. SMC-Ca^2+^: ***p=0.0001, Vasomotor dynamics vs. PVF-Ca^2+^: **p=0.0026, SMC-Ca^2+^ vs. LPMF-Ca^2+^: **p=0.0025. Arteriolar ROIs n=32, SMC-Ca^2+^ ROIs n=25, PVF-Ca^2+^ ROIs n=55, LPMF-Ca^2+^ ROIs n=130 in 3 Col1a2CreER-GCaMP6f mice. Each point is an individual cell. **(h)** Graph showing phase-locking values of SMC-, PVF-, and LPMF-Ca^2+^ oscillations with vasomotor oscillations. Same n values as shown in (g). One-way ANOVA followed by Tukey’s multiple comparison test: SMCs vs. LPMFs: ****p<0.0008. Cells with a PLV ≥0.2 (above dashed line) were included in downstream analyses shown in (m) and (n). **(i)** Histograms displaying phase relationships reported as circular means (°) for percent (%) of SMC-, PVF-, and LPMF-Ca^2+^ ROIs with vasomotor oscillations. Same n values as shown in (g). Cells grouped into 30° bins from -180 to +180 from peak dilation. Mean for each cell type and peak dilation indicated with dashed lines. Annotations provided to assist in interpretation of preferred phase relationship for cellular-Ca^2+^ before, at, or after diameter peak. **(j)** Graphs showing 30 seconds of % vessel diameter changes (black) with one SMC-Ca^2+^ (red) example, two PVF-Ca^2+^ (purple) examples, and two LPMF-Ca^2+^ (blue) examples with their respective **(j’)** cross-correlation graphs below indicating their strongest correlation coefficient (r), timing relationship (t), and whether Ca^2+^ or diameter leads. **(k)** Graph of peak correlation of cellular-Ca^2+^ and diameter oscillations following cross-correlation analysis. Same n values as shown in (g). PVF-Ca^2+^ ROIs: 27 positive-dominant (pink), 28 negative-dominant (green); LPMF-Ca^2+^ ROIs: 69 positive-dominant, 61 negative-dominant. Pearson correlation coefficient, time lag, and leading signal indicated on each graph. **(l)** Bar graph showing the proportion of positive-dominant and negative-dominant SMCs, PVFs and LPMF based on their peak correlation with Ca^2+^ and vasomotor dynamics. Same n values as shown in (g). SMC-Ca^2+^ ROIs: 6 positive-dom., 19 negative-dom. **(m)** Graphs of cycle-triggered average of cellular-Ca^2+^ and arteriolar dynamics for SMCs (red), positive-dominant (pink) and negative-dominant (green) PVFs and LPMFs. Dynamics of Ca^2+^ and diameter are shown as a normalized change over the mean. Cells with a PLV ≥0.2 (above dashed line in (h)) were included in downstream analyses shown in (n). SEM indicated for mean Ca^2+^ and diameter. **(n)** Graphs showing the timing of the closest Ca^2+^ troughs and peaks from peak vessel dilation centered around 0 seconds for SMCs, PVFs, and LPMFs. One-way ANOVA followed by Kruskal-Wallis test: Closest Ca^2+^ trough vs. peak - SMCs: ****p<0.0001, PVFs: *p=0.0135. SMCs vs. LPMFs - Closest Ca^2+^ peak: **p=0.0026. **(o)** Graph modeling the dynamic range of (left graph) PVF-Ca^2+^ oscillations (purple) with SMC-Ca^2+^ (red) and (right graph) LPMFs (blue) during a vasomotor cycle (black). Range determined by collating positive- and negative-dominant variation for PVFs and LPMFs shown in (m).

To determine whether PVFs participate in the vasomotor dynamics of pial arterioles *in vivo*, we performed awake *in vivo* imaging on Col1a2CreER-GCaMP6f mice during spontaneous dilation and constriction events (**Fig. 1e**). We measured Ca^2+^ activity in PVFs, SMCs, and LPMFs alongside vasomotion (**Fig. 1f**). SMC- and PVF-Ca^2+^ oscillations occurred at significantly higher frequencies than vasomotion, whereas LPMF-Ca^2+^ oscillations were slower than SMC-Ca^2+^ oscillations but not significantly different from vasomotion (**Fig. 1g**). Phase analysis revealed that SMCs were phase-locked with vasomotor dynamics and exhibited constrained phase angles, indicating consistent coupling with the vasomotor cycle (**Fig. 1h, i and Extended Data Fig. 3a**). PVFs displayed slightly broader phase relationships while maintaining phase-locking comparable to SMCs, suggesting heterogeneous Ca^2+^ activity within a vasomotor cycle. LPMFs exhibited reduced phase-locking and more diffuse phase distribution indicating a weaker temporal organization with vasomotor oscillations. These data, together with Rayleigh tests and von Mises concentration, indicate PVF-Ca^2+^ oscillations, unlike LPMF-Ca^2+^, have clustered, non-random phase relationships with vasomotion (**Extended Data Fig. 3b-d**).

To resolve how PVF-Ca^2+^ changes relate to fluctuations in vessel diameter, we performed cross-correlation analyses on SMC-, PVF-, and LPMF-Ca^2+^ activity with vessel diameter (**Fig. 1j, j’**). SMC-Ca^2+^ displayed a predominant inverse relationship with arteriolar diameter, whereas PVF- and LPMF-Ca^2+^ segregated into positive-dominant and negative-dominant relationships based on their peak correlation with vasomotion. Some SMCs displayed positive peak correlations which we address in subsequent analyses (**Extended Data Fig. 4d**). Peak correlation coefficients were comparable among SMCs and negative-dominant PVFs and LPMFs, as were those of the positive-dominant PVFs and LPMFs (**Fig. 1k**). PVFs and LPMFs were relatively evenly distributed between the positive- and negative-dominant populations, with a small population classified as uncorrelated (r ≤ ±0.1)(**Fig. 1l**). Frequency, phase-locking, and phase relationships were similar between SMCs and PVFs classified as positive-dominant and negative-dominant (**Extended Data Fig. 3e-g**). However, LPMF subpopulations were consistently different from SMCs.

Cycle-triggered averaging on cells with phase-locking values ≥0.2, representing cells with at least intermittent phase-locking, revealed that SMC-Ca^2+^ decreased before peak arteriole dilation (**Fig. 1n, m; Extended Data Fig. 4a-c**). A minority of SMCs exhibited positive peak correlations with vessel diameter. However, these cells exhibited phase angles shifted modestly toward peak dilation but retained Ca^2+^ troughs and peaks similarly to negatively-correlated SMCs, suggesting that correlation sign reflects differences in temporal positioning within the vasomotor cycle rather than fundamentally distinct Ca^2+^ behaviors (**Extended Data Fig. 4d**). Negative-dominant PVFs exhibited similar temporal relationships to the vasomotor cycle as SMCs, whereas positive-dominant PVFs exhibited Ca^2+^ increases that more closely followed peak dilation. LPMFs, both positive- and negative-dominant groups, did not exhibit Ca^2+^ patterns that closely followed the vasomotor cycle. Together, these findings indicate that PVF-Ca^2+^, unlike LPMF-Ca^2+^, is temporally coupled to vasomotion but exhibit broader timing relationships than SMCs (**Fig. 1o**).

To determine whether PVFs respond during functional hyperemia, we performed repeated whisker stimulation experiments on awake Col1a2CreER-GCaMP6f mice while imaging over the barrel cortex (**Extended Data Fig. 5a**). Across repeated whisker stimulations (**Extended Data Fig. 5b**), approximately 50% of the PVFs maintained persistent increase or decrease Ca^2+^ responses across repeated trials, whereas the remainder switched between response states indicating dynamic adaptation of PVF-Ca^2+^ signaling (**Extended Data Fig. 5c**). Overall, ∼65% of PVF responses were associated with a decrease in their Ca^2+^ levels, ∼25% increase in Ca^2+^, and ∼9% were non-responsive (**Extended Data Fig. 5d**). Increases in PVF-Ca^2+^ preceded SMC responses and vessel dilation whereas decreases in PVF-Ca^2+^ occurred slightly after SMC-Ca^2+^ decreases (**Extended Data Fig. 5e-h**). Together, these data indicate that PVFs exhibit dynamic Ca^2+^ responses during functional hyperemia, with a majority of PVFs decreasing Ca^2+^ along with SMCs^14^.

To determine whether PVFs can influence vasomotor dynamics, we generated Col1a2CreER-GqDREADD mice in which tamoxifen administration induces fibroblast-specific expression of hM3Dq, the excitatory Gq-coupled designer receptor exclusively activated by designer drugs (DREADDs). Recombination was verified by mCitrine which is co-expressed with hM3Dq in this mouse line^15^ (**Fig. 2a**). Following intraperitoneal administration of Deschloroclozapine (DCZ), a brain penetrant agonist for DREADDs, which activates PLC/IP3 signaling mobilizing intracellular Ca^2+^ stores^16^, we observed a sustained increase in pial arteriole diameter from 10-60 mins post DCZ. Critically, dilation was localized to vessel segments containing mCitrine-positive PVFs, suggesting a highly local effect (**Fig. 2a-c**). In contrast, vessel segments lacking mCitrine-positive PVFs (mCitrine-negative) and littermate controls showed little change in vessel diameter following DCZ. Although Col1a2CreER recombines SMCs, we did not detect mCitrine expression in SMCs in Col1a2CreER-GqDREADD mice, potentially reflecting differences in reporter expression associated with the GqDREADD allele compared with Rosa26-targeted reporter lines such as Ai14 or Ai95.

**Figure 2.**
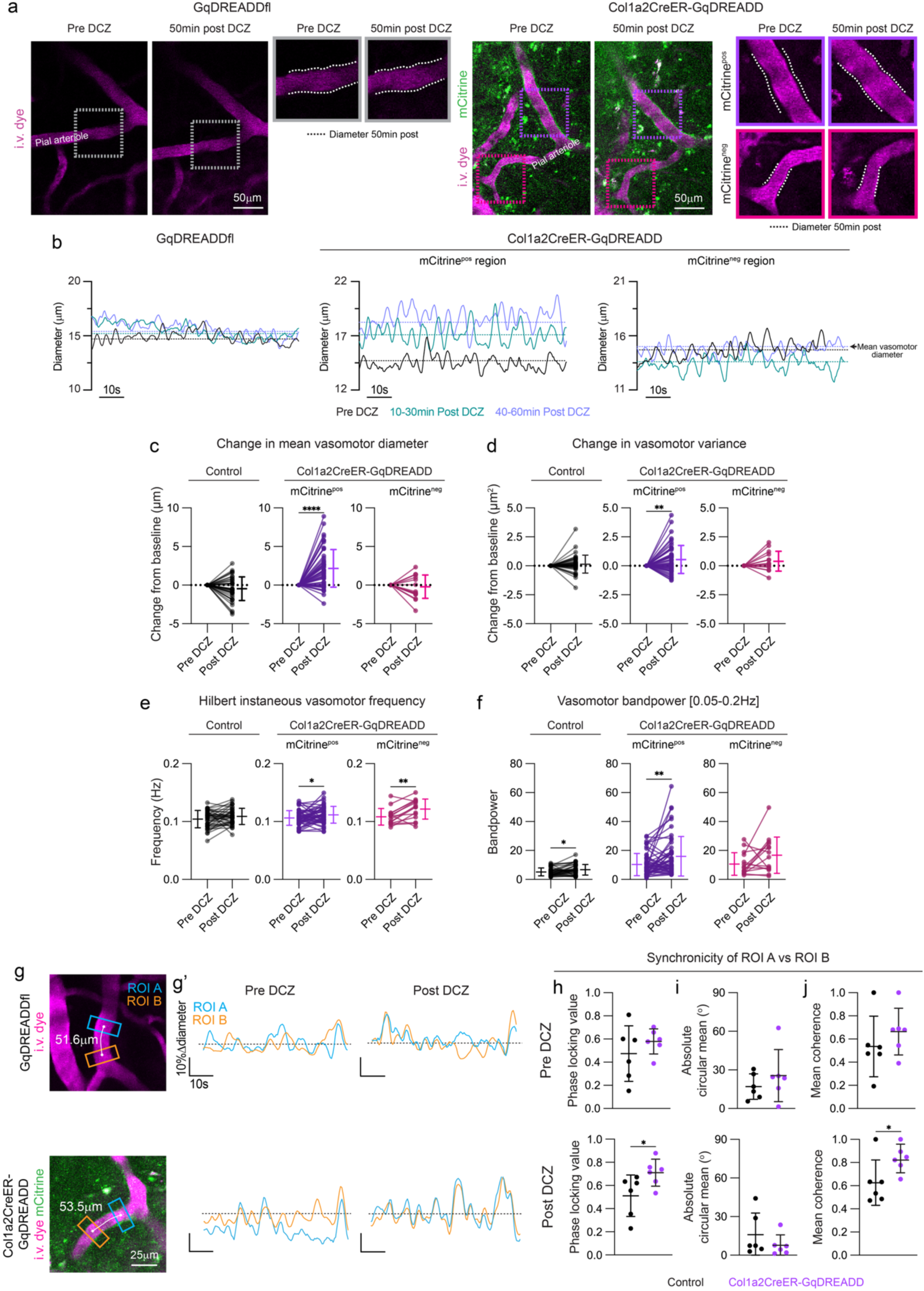
Activation of perivascular fibroblast-Gq signaling enhances arteriolar vasodynamics. **(a)** Representative *in vivo* images of pial arterioles (magenta; i.v. dye; 70kDa Texas Red-Dextran) from a GqDREADDflox (GqDREADfl) littermate control and Col1a2CreER-GqDREADD mice pre-Deschloroclozapine (DCZ) and 50 mins post DCZ. Diameter region of interests shown in respective insets for control (GqDREADDfl) as well as mCitrine-positive (mCitrine^pos^; green fluorescence) and mCitrine-negative (mCitrine^neg^) vascular territories. Diameter 50min post DCZ outlined with white dashed lines overlaid on pre DCZ to demonstrate diameter change in mCitrine^pos^ region. **(b)** Graphs showing 1 min of resting diameter changes in the vascular territories from vessel regions shown above from the GqDREADDfl mouse and mCitrine^pos^ and mCitrine^neg^ regions in the Col1a2CreER-GqDREADD mouse. Diameter changes overlaid for pre DCZ (black), 10-30 min post DCZ (green), and 40-60 min DCZ (purple) for each respective vascular territory shown in (a). Mean vasomotor diameter pre and post DCZ timeframes indicated by dashed lines. **(c-f)** Graphs showing the **(c)** change in vasomotor diameter mean, **(d)** change in vasomotor variance, **(e)** Hilbert instantaneous frequency, and **(f)** vasomotor bandpower within 0.05-0.2 Hz pre-DCZ and post-DCZ (40-60 min post) for ROIs along pial arterioles in control (black; GqDREADDfl or Col1a2CreER) as well as mCitrine^pos^ (purple) and mCitrine^neg^ (pink) regions in Col1a2CreER-GqDREADD mice. Col1a2CreER-GqDREADD (n=5 mice) mCitrine^pos^ arteriolar ROIs n=51. Paired t-test: pre-vs. post-DCZ; change in vasomotor diameter mean ****p<0.0001, change in vasomotor variance **p=0.0021, change in vasomotor frequency *p=0.0204, change in vasomotor bandpower **p=0.0028. mCitrine^neg^ arteriolar ROIs n=14: Paired t-test; pre- and post-DCZ; change in vasomotor frequency **p=0.0034. Control (Col1a2CreER or GqDREADDfl; n=3 mice). Paired t-test: pre- and post-DCZ; change in vasomotor bandpower *p=0.015. **(g)** Representative images of analysis for vessel synchronicity in GqDREADDfl and Col1a2CreER-GqDREADD mice showing the mapping of ROI A (blue) and ROI B (orange) with their respective vascular distances around 50µm. **(g’)** Graphs showing 1 min of resting diameter changes in ROI A and ROI B as shown in (g) pre- and post-DCZ. **(h-j)** Graphs showing the synchronicity of ROI A vs ROI B via **(h)** phase-locking value, **(i)** absolute circular mean (°), and **(j)** mean coherence in control (black) and Col1a2CreER-GqDREADD (purple) mice. Cola2CreER-GqDREADD: n=6 pial arterioles, 3 mice and Control: n=6 pial arterioles; 3 mice. Pre-DCZ analysis: Col1a2CreER-GqDREADD vs control: Welch’s t-test: phase-locking value p=0.36, Mardia-Watson-Wheeler test: absolute circular mean (°) W=1.22, df=2, p=0.543, mean coherence p=0.36. Post-DCZ analysis: Col1a2CreER-GqDREADD vs control: Welch’s t-test: phase-locking value post DCZ *p=0.049. Mardia-Watson-Wheeler test: absolute circular mean (°) W=0.164, df=2, p=0.921, mean coherence *p=0.043.

This increase in vessel diameter was accompanied by enhanced vasomotor dynamics within mCitrine-positive segments, as evidenced by increased diameter variance (**Fig. 2d and Extended Data Fig. 6**), while variance remained stable in mCitrine-negative segments and control mice. Although vasomotor frequency remained unchanged in control mice, mCitrine-positive and negative segments exhibited increases in vasomotor frequency post DCZ (**Fig. 2e**). DCZ modestly increased vasomotor bandpower in control mice (**Fig. 2f**), indicating a limited effect on vasodynamics. In contrast, fibroblast-Gq activation increased the spectral power of diameter oscillations within the vasomotor frequency band (0.05-0.25Hz) along mCitrine-positive segments. Together, the concurrent increases in vessel diameter, variance, and vasomotor bandpower within mCitrine-positive segments demonstrate that PVFs can locally modulate vasomotor state.

We next examined whether PVFs influence vasomotor coordination. We quantified vasomotor dynamics of two comparably spaced ROIs (∼50µm; ROI A and ROI B) positioned along continuous, non-branching arteriole segments in control and Col1a2CreER-GqDREADD mice (**Fig. 2g and Extended Data Fig. 7**). Analyses in Col1a2CreER-GqDREADD mice were restricted to vessel segments with continuous mCitrine-positive coverage. Prior to DCZ administration, phase-locking, absolute circular means, and mean coherence were comparable between genotypes (**Fig. 2h-j and Extended Data Fig. 8**). Following fibroblast-Gq activation, Col1a2CreER-GqDREADD mice demonstrated a significant increase in both phase-locking and mean coherence compared with controls. Together, these findings indicate that enhanced PVF-Gq signaling promotes a locally coordinated vasomotor state characterized by increased oscillatory activity and improved spatiotemporal organization of vasomotion.

We next examined whether enhanced fibroblast-Gq signaling altered tracer-labeled CSF influx 30 mins post DCZ exposure using intracisternal magna injections of 45 kDa Ova-647 and 10 kDa lysine-fixable FITC (**Fig. 3a**). Quantification of tracer distribution along the cortical surface and penetrating arterioles (**Fig. 3b,c**), revealed increased tracer coverage along both compartments in Col1a2CreER-GqDREADD mice, including a trend toward increased cortical depths (**Fig. 3d-g**). OVA-647 and FITC displayed similar distribution, and the enhancement of CSF tracer influx in Col1a2CreER-GqDREADD mice was maintained along the rostral-caudal axis (**Extended Data Fig. 9**). These findings suggest that enhanced vasomotor dynamics through chemogenetic activation of fibroblast-Gq signaling facilitates increased perivascular fluid influx.

**Figure 3.**
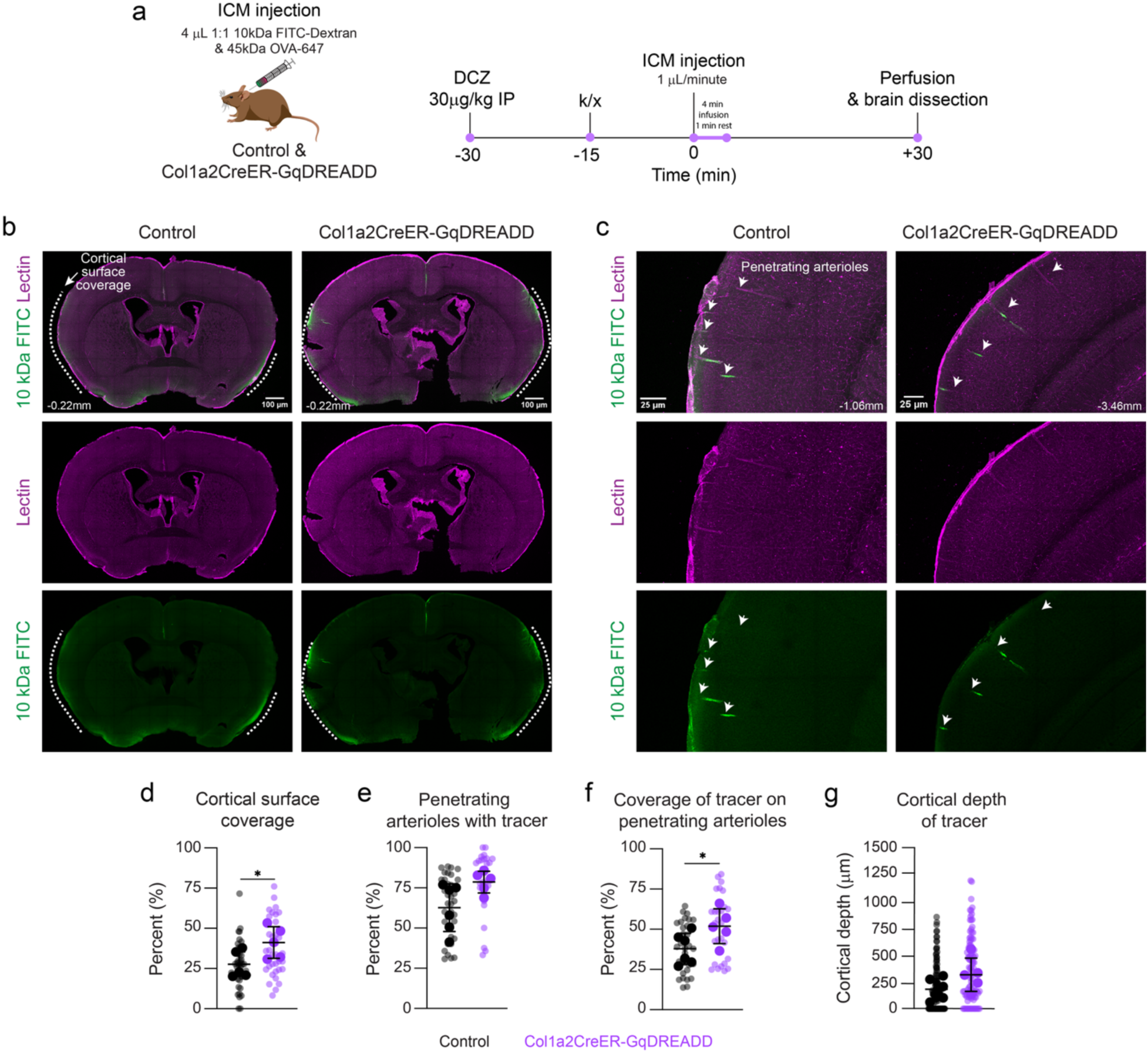
Activation of Gq signaling in perivascular fibroblasts accelerates peri-arteriolar CSF influx. **(a)** Graphical depiction of intracisterna magna injection (ICM) and experimental timeline on Col1a2CreER-GqDREADD and control mice wherein mice received Deschloroclozapine (DCZ; 30µg/kg) via intraperitoneal injection 30 minutes prior to ICM injection. Fifteen minutes prior to ICM injection, mice were anesthetized with ketamine (105mg/kg)/xylazine (15mg/kg). Mice received 4 µL of 1:1 mix of 5 mg/mL 10 kDa FITC-dextran (lysine-fixable) and 1 mg/mL 45 kDa OVA-647 infused at a rate of 1 µL/min (4 min total) followed by a 1 min waiting period. Thirty minutes following ICM injections, mice were then prepared for PFA-perfusions and dissections. **(b & c)** Confocal images of **(b)** whole brain sections of cortical surface tracer distribution (white dashed line) and **(c)** zoomed in images of cortical penetrating arterioles (white arrows) with peri-arteriolar tracer in controls and Col1a2CreER-GqDREADD mice. Lectin depicted in magenta, 10 kDa FITC Dextran shown in green. Corresponding positions relative to bregma noted. **(d-g)** Graphs of **(d)** Cortical surface coverage, **(e)** Percent penetrating arterioles with tracer, **(f)** Coverage of tracer on penetrating arterioles, and **(g)** Cortical depth of tracer in control (n=5 mice) and Col1a2CreER-GqDREADD (n=5 mice). Unpaired t-test: Cortical surface coverage: control vs Col1a2CreER-GqDREADD: *p=0.0296. Penetrating arterioles with tracer: p=0.0543. Coverage of tracer on penetrating arterioles: *p=0.491. Cortical depth of tracer: p=0.1362. Small points indicate individual tissue sections from each mouse, larger points denote the mean value for each mouse. Statistics were performed on mice, with quantification from individual sections shown for visualization only.

Here, we identify a role for brain PVFs in influencing the vasodynamics of pial arterioles, suggesting these cells are a novel control point in the regulation of spontaneous arteriolar oscillations. The temporal relationship between PVF-Ca^2+^ dynamics and vessel diameter raises the possibility that PVFs monitor ongoing vasomotor activity and are sensitive to mechanical cues arising from cyclic diameter changes and/or SMC contractility. How enhanced Gq signaling in PVFs locally augments vasomotion remains unknown and warrants further investigation. Potential mechanisms include Ca^2+^-dependent signaling pathways, modification of vascular biomechanics, or communication with neighboring SMCs. Recent work showing that PVFs regulate vasodilation in the corpus cavernosa supports their emerging roles in vascular physiology^17^. A limitation of our studies is that Col1a2CreER targets fibroblasts throughout the body, as no brain-specific PVF Cre line is currently available, raising the possibility that peripheral fibroblast activation may contribute to effects on CSF influx. Finally, impaired vasodynamics and perivascular fluid transport are becoming increasingly associated with brain waste accumulation in aging and neurodegenerative diseases^5^. Our findings suggest that PVFs may support vascular-related brain waste clearance mechanisms and that pathological changes in PVF populations may contribute to impaired vasomotor dynamics and waste accumulation in disease^18–20^.

## Methods

### Animals

Mice were housed in a specific pathogen-free facility approved by AAALAC and were handled in accordance with protocols approved by IACUCs of University of Colorado Anschutz Medical Campus (CU Anschutz) and Seattle Children’s Research Institute (SCRI). Col1a2CreER-iDTRflox, Col1a2CreER-Ai14flox, Col1a2CreER-Ai95flox and Col1a2CreER-GqDREADDflox mouse lines were created by breeding Col1a2CreER mice (Jax #029567) with ROSA26iDTR-flox (Jax #007900), Ai14flox (Jax #007914), Ai95-flox (Jax #028865), and GqDREADD-flox (Jax #026220). Experimental mice included were of the following genotypes: Col1a2CreER/+; iDTRfl/+ (Col1a2CreER-iDTR), Col1a2CreER/+; Ai14fl/+ (Col1a2CreER-tdTomato), Col1a2CreER/+; Ai95fl/+ (Col1a2CreER-GCaMP6f), and Col1a2CreER/+; GqDREADDfl/+ (Col1a2CreER-GqDREADD). Littermate control mice (Col1a2CreER/+, iDTRfl/+, or GqDREADDfl/+) were used where necessary. Col1a1GFP mice, created by David Brenner^21^, were used for morphological characterization of PVFs in *ex vivo* arteriolar preparations. To induce CreER recombination in brain fibroblasts, mice were given 1 to 2 days of tamoxifen (80mg/kg i.p dissolved in corn oil). This minimized recombination in SMCs to ∼4% of the perivascular cells^4^. Both male and female mice within 3 to 12 months of age were utilized.

### Cranial window surgeries

In Col1a2CreER-GCaMP6f, Col1a2CreER-GqDREADD mice and their littermate controls (GqDREADDfl/+ or Col1a2CreER/+), to minimize perturbation of the brain, 3-mm diameter circular thin-skull cranial windows were placed over the somatosensory cortex for *in vivo* imaging, as previously described with the exception of window polishing due to the acute nature of these experiments. Following these surgeries, mice recovered for 1-2 days prior to *in vivo* imaging. For Col1a2CreER-iDTR mice and their respective littermate controls (iDTRfl/+ or Col1a2CreER/+), 3-mm diameter skull-removed cranial windows were created to allow for topical application of diphtheria toxin and AAV8-CAG-GFP to the surface of the brain and subsequent depletion of brain fibroblasts and is described in more detail below. Skull-removed cranial windows were also implanted in Col1a2CreER-tdTomato mice. Extra soaking with artificial cerebrospinal fluid (aCSF), which included at least 3 separate 1 min pauses with aCSF, during drilling ensured softening of the bone and separation of the dura from the overlying skull during removal^4^.

### Diphtheria-mediated fibroblast depletion experiments

Following removal of a 3-mm skull segment overlying the somatosensory cortex, bleeding was controlled using Surgifoam (Ethicon) soaked in aCSF. A solution containing 1 ng diphtheria toxin (DTX; Sigma, D0564) and 10^12^ genome copies of AAV8-CAG-GFP^10^ (Addgene, 37825-AAV8) in 30 μL aCSF was applied directly to the cortical surface and allowed to remain undisturbed for at least 3 min. A cranial window coverslip plug, composed of 4 mm and 3 mm round coverglasses secured with UV-curable optical glue, was then secured in place with Loctite 401 super glue and dental cement to seal the DTX and AAV solution. Three days following DTX/AAV exposure, GFP expression was present as a thin sheet around mainly pial arterioles indicative of PVFs and appeared punctate in the meningeal fibroblast populations suggesting ongoing transgene expression and spread.

### Awake in vivo imaging

To label and image the brain vasculature, fluorescent dextrans were injected retro-orbitally under deep isoflurane anesthesia (2% MAC in medical-grade air). All mice were injected with 25 µL of 2.5% (w/v in saline) 70 kDa Texan Red-dextran (Invitrogen; D1830), or 25 µL of 5% (w/v in saline) custom Alexa Fluor 680 (Life Technologies; A20008) conjugated to 2MDa Dextran (Fisher Scientific; NC1275021). All mice were habituated the day prior to imaging experiments which entailed 20-30 min head fixation on a treadmill (PhenoSys speedbelt at SCRI or Low-friction compact treadmill for mice at CU Anschutz) which allowed free forward-backward movement. The following subsequent days, mice underwent awake *in vivo* imaging for up to 2 hours with collection of images and movies occurring no earlier than 10 minutes following isoflurane exposure for the retro-orbital injections. Imaging was performed with a Bruker Investigator coupled to a Spectra-Physics Insight X3 (SCRI) or MaiTai (CU Anschutz) laser. The laser was tuned to 920 nm excitation for all experiments. Collection of green, red, and far-red fluorescence emission was achieved with 525/70, 595/50, and 660/40 emission bandpass filters respectively, and detected with Hamamatsu GaAsP photomultiplier tubes. A 20x (1.0 NA) water-immersion objective (Olympus; XLUMPLFLN) was used to collect high-resolution images. For higher resolution imaging of brain fibroblasts and the vasculature, z-stacks were collected at 1.0 μm z increments with a 347 μm x 347 μm (512 × 512 pixel resolution) field-of-view using 3.6 μs/pixel dwell time. For imaging of Ca^2+^ signaling and vasomotor dynamics in Col1a2CreER-Ai95flox mice, movies were captured at 2 different frequencies: 1) 1.951 Hz with a 295 μm × 295 μm (256 × 256 pixel resolution) field-of-view averaging every two frames using 1.2 μs/pixel dwell time for 307.2 s, or 2) 7.44Hz using galvo resonant imaging with a 196 μm x 196 μm (512 x 512 pixel resolution) field-of-view averaging every four frames for 179.8s. The use of these two imaging speeds allowed us to determine the best timing relationship for brain fibroblast-Ca^2+^ signaling and vasomotion. For imaging of vasomotion in Col1a2CreER-iDTRflox and Col1a2CreER-GqDREADD mice were imaged at 1.951 Hz.

### Parenchymal arteriole preparation

Mice were euthanized by intraperitoneal administration of sodium pentobarbital (100 mg/kg), followed immediately by decapitation. Cortical parenchymal arterioles (PAs) were isolated as previously described^12^ with emphasis on superficial PAs. Briefly, brains were rapidly removed and immersed in ice-cold MOPS-buffered saline containing 135 mM NaCl, 5 mM KCl, 1 mM KH_2_PO_4_, 1 mM MgSO_4_, 2.5 mM CaCl_2_, 5 mM glucose, 3 mM MOPS, 0.02 mM EDTA, 2 mM pyruvate, and 10 mg/mL bovine serum albumin (BSA) (pH 7.3). Under a dissecting microscope, segments of the middle cerebral artery (MCA) with the surrounding cortical tissue were carefully dissected from both hemispheres and carefully dissected free from the surrounding neural tissue. PAs were cannulated and tied off while the other end was occluded as previously described^22^. The preparation was continuously superfused with oxygenated aCSF at 4 mL/min and maintained at 36.5 ± 1°C. The aCSF was equilibrated with a gas mixture of 5% CO_2_, 20% O_2_, and 75% N_2_ and contained (in mM): 125 NaCl, 3 KCl, 26 NaHCO₃, 1.25 NaH_2_PO_4_, 1 MgCl_2_, 4 glucose, and 2 CaCl_2_ (pH 7.3).

### Morphological distinction between perivascular fibroblasts and smooth muscle cells

We captured confocal images of isolated pial arteries and penetrating arterioles from Col1a1GFP mice to verify the preservation of PVF populations in *ex vivo* vessel preparations. These images were also used to compare PVF morphology between Col1a1GFP and Col1a2CreER-GCaMP6f mice for *ex vivo* PA experiments. In addition, we performed *in vivo* imaging on Col1a2CreER-tdTomato mice to compare the morphological characteristics of tdTomato-expressing SMCs and PVFs with GCaMP6f-expressing SMCs and PVFs in the Col1a2CreER-GCaMP6f mouse line. To categorize PVFs and SMCs in our *in vivo* and *ex vivo* Ca^2+^ experiments as shown in **Extended Data Fig. 2**, PVFs were classified by their flattened somata and thin, fibrillar processes extending along the outside of pial arteries and penetrating arterioles. SMCs were classified by circumferentially-oriented morphology and characteristic banding pattern due to their wrapping around the arterial wall. GCaMP6f-positive perivascular cells exhibiting overlapping or insufficient morphological features to classify as PVF or SMC along the vessel wall were not classified and excluded from analyses in these studies. In the *ex vivo* preparations from Col1a2CreER-GCaMP6f mice, PVFs were very bright under basal conditions (aCSF bath) whereas recombined SMCs expressing GCaMP6f were dimmer basally and became readily visible upon 60mM K^+^-induced vasoconstriction and the subsequent increase in SMC-Ca^2+^.

### Calcium imaging of perivascular fibroblasts and smooth muscle cells ex vivo

Post parenchymal arteriole preparation, the vessel organ chamber was placed on a Nikon A1R Ti2 inverted confocal microscope using a CFI60 Plan Fluor 20X water immersion objective lens (NA 0.95, WD 0.95 mm, FOV 22 mm, DIC). Single-plane t-series images were focused on GCaMP6f-expressing PVFs under basal conditions (aCSF), bath application of 60mM K^+^, and increases in intraluminal pressure (20 mmHg to 100 mmHg) to capture Ca^2+^ (GCaMP6f; *λ*Ex: 488 nm; *λ*Em: 525 nm) events at 3.67 frames per second, via Galvano scanner. Under basal conditions, GCaMP6f-expressing PVFs were visible whereas 60mM K^+^ rapidly increase GCaMP6f fluorescence in SMCs which were distinguished by their banding morphology demonstrated in **Extended Data Fig. 2**. Two to three PVFs and SMCs were selected per vessel. Regions of interests (ROIs) were analyzed using a custom-designed noncommercially available SparkAn software (Dr. Adrian Bonev, University of Vermont). F/F_0_ was normalized by taking baseline or basal Ca^2+^ fluorescence of 10 images. Maximum F/F_0_ peaks were selected for summary data^13^. Vessels that did not respond to 60mM K^+^ were excluded. Vessel diameter was measured offline using VasoTracker^23^. Image sequences were analyzed by placing line-scan ROIs perpendicular to the vessel walls at defined locations along the arteriole. VasoTracker automatically detected vessel edges and calculated luminal diameter over time. All measurements were visually inspected to ensure accurate edge detection, and diameter traces were exported into GraphPad Prism for further analysis. Changes in vessel diameter were calculated relative to baseline values acquired immediately before experimental interventions using the following calculation: *((Diameter_passive_ – Diameter_active_)/Diameter_passive_) *100*.

### Analysis of in vivo pial arteriole vasomotor and cellular-Ca^2+^ dynamics

Movies were captured of the brain surface encompassing the meningeal surface and pial arterioles. For each imaging session, the longest contiguous resting period of at least 60 s was selected for analysis. Resting periods were identified directly from the movies by the absence of movement-related displacement of the imaging plane. In contrast, active treadmill locomotion produced obvious motion artifacts and concomitant vasodilation. Locomotion-induced movement was readily detectable, allowing these periods to be reliably excluded from analysis. To analyze vasomotion, the VasoMetrics plugin was utilized to measure vessel diameter in each frame. To analyze cellular-Ca^2+^ signatures in FIJI, an ROI was drawn around each cell soma. ROIs were always ensured to encompass the target cells across all movie frames. In the event a major shift was observed, ROIs were redrawn for those portions of the movies to always maintain consistency in where the GCaMP6f signal was being measured. The fluorescent intensity profile throughout each movie was obtained using the plot z-axis profile function and the change in fluorescence divided by the mean fluorescence intensity (ΔF/F_0_) was calculated for the longest contiguous resting period. Analysis of vasomotor dynamics in Col1a2CreER-GqDREADD and Col1a2CreER-iDTRflox mice (and littermate controls) was done by selecting ROIs along the pial arterioles with at least 20 µm of vascular length separation and measuring vessel diameter in the movies to identify local diameter changes along pial arterioles.

Downstream analyses, including mean, variance (square of the standard deviation), frequency, bandpower, cross-correlation, Hilbert transform and phase analyses, cycle-triggered averaging, and mean coherence, were all performed on the longest contiguous resting period for each ROI. The Hilbert instantaneous frequency of resting Ca^2+^ and diameter were calculated in MATLAB by first using the hilbert() function to extract instantaneous phase from the signals, and instantaneous frequency was subsequently calculated from the temporal derivative of the unwrapped instantaneous phase, multiplied by the sampling frequency and divided by 2π. Because vasomotor and cellular Ca^2+^ oscillations exhibit cycle-to-cycle variability and recordings were limited to relatively short resting epochs, we used Hilbert instantaneous frequency to quantify the frequency of each oscillatory cycle rather than relying on a single mean frequency estimate. Power spectral density and bandpower using Welch’s method were calculated using pwelch() and bandpower() MATLAB functions. For frequency and spectral power analyses, signals were bandpass filtered between 0.05-0.2Hz to isolate the canonical vasomotor frequency range and minimize contributions from slower fluctuations. This lies within the frequency band known to represent vasomotor activity^7^ (0.025-0.25Hz). Peak correlation and classification of PVFs and LPMFs as positive or negative was identified as ROIs with r values <0.1 or >0.1; values between -0.1 and 0.1 were classified as uncorrelated. To characterize the temporal relationship between cellular Ca^2+^ signaling and arteriolar vasodynamics throughout recurring oscillatory cycles over time, the instantaneous phase was derived via the Hilbert transform. Ca^2+^ and vasomotor signals were bandpass filtered using second-order zero-phase Butterworth filtering using filtfilt() and butter() MATLAB functions for frequencies in the 0.025–0.25 Hz range, which preserved slower vasomotor cycles and maintained waveform continuity required for stable instantaneous phase estimation. Hilbert transform was performed on filtered signals using the hilbert() MATLAB function. Circular mean phase difference (̅Δϕ) was calculated from the distribution of instantaneous phase differences as the circular mean of the phase offsets using the following equation: *̅Δϕ=atan2(⟨sin(Δϕ)⟩,⟨cos(Δϕ)⟩* where *Δϕ* is the instantaneous phase difference and *⟨⋅⟩* denotes the arithmetic mean across observations. Phase-locking value (PLV) was calculated as the length of the mean resultant vector of the instantaneous phase differences: *PLV* = |(1/*N*) *Σi*₌₁ᴺ *e*ⁱ*Δφi*|, where *N* = total number of observations (time points), *k* = index of each observation (*k*=1,…,*N*), *e*= the base of the natural logarithm, *i* = √−1 is the imaginary unit, and *Δϕ_i_* = instantaneous phase difference between the cellular Ca^2+^ signal and the vasomotor oscillation at observation *i*. The Rayleigh test for uniformity was performed to determine whether instantaneous phase differences displayed significant directional clustering rather than a uniform distribution. Rayleigh’s test statistic (*Z=nR^−2^*) was also computed, where *n* is the number of observations and *R̄* is the mean resultant length. The von Mises concentration (κ), which quantifies the concentration of phase differences around the mean phase, was estimated from the mean resultant length (*R̄)* using standard circular statistical methods. To further characterize the timing of cellular Ca^2+^ activity relative to vasomotor cycles, cycle-triggered averaging was used by aligning Ca^2+^ oscillations to oscillatory vasomotor peaks, identified as alignment points using the findpeaks() function in MATLAB. Ca^2+^ and diameter signals were again bandpass filtered using second-order zero-phase Butterworth filtering using filtfilt() and butter() functions. Mean cycle-triggered traces and SEM were calculated across cycles.

### Cortical whole mount dissection, staining and analysis

Cortical whole mounts, and subsequent immunostaining and imaging of pial vessels, were prepared from the Col1a2Cre-iDTR and their respective littermate controls (iDTRflox) with cranial windows 7-days post DTX exposure. Mice were anesthetized with euthasol and subsequently underwent trans-cardial perfusions with PBS followed by 4% paraformaldehyde. Brains were then dissected and post-fixed for 24 hours followed by preparation for dissection to isolate the pial surface for immunostaining in PBS with 1% sodium azide. Brains were then hemisected and the superficial layers of the cortex were dissected away from the underlying corpus callosum using a scalpel to obtain the cortical surface surrounding the cranial window territory. Cortical whole mounts were then immunostained for anti-ɑSMA-Cy3 (Millipore Sigma; C6198) to label SMCs and anti-GFP-488 (ThermoFisher; A-21311) to label AAV8-CAG-GFP-labeled fibroblasts in a solution of 2% TritonX-100, 5% BSA and 0.1% sodium azide in PBS for 48 hours at 4°C. Whole mounts were then washed 3 times for 5 minutes in PBS and mounted with Fluromount-G (ThermoFisher; 00-4958-02) onto 0.9mm cover slip wells (Millipore Sigma; GBL635011). Three immunofluorescent images per animal were captured with a 20x objective using a Zeiss 780 LSM confocal microscope. Percent ɑSMA coverage (ɑSMA length/total pial arteriole length x 100) was quantified by measuring ɑSMA length and total pial arteriole length using FIJI line segment tool.

### Whisker stimulation and analysis in awake mice

During awake imaging experiments, Col1a2CreER-Ai95flox mice were head-fixed on a treadmill and received retro-orbital dye injections of i.v. dye mice under isoflurane anesthesia (2% MAC in medical-grade air). Ten minutes after the mouse recovered from isoflurane anesthesia, multiple sites of pial arterioles supplying mainly the barrel-cortex on the lateral side of the window were targeted for whisker stimulation experiments. Three separate movies were captured per area at 1.951 Hz spanning two minutes with 10 s Picospritzer-driven air puffs at 8 Hz after 60 s followed by 50 s of rest prior to the next whisker stimulation trial. ROIs for cellular-Ca^2+^ were selected and distinguished for PVFs and SMCs based on the banded morphology of SMCs and flattened soma of PVFs and fluorescent intensity was measured as described above. Their respective vascular territories were measured for diameter changes throughout all three whisker stimulation trials using Vasometrics. Time to peak following the whisker stimulation was identified per ROI, including arterial dilation, as well as an immediate increase or decrease in cellular-Ca^2+^ where Ca^2+^ responses less than 5% change in ΔF/F0 were classified as non-responsive. This allowed us to classify cells as either “positive”, “negative”, or “non-responsive” PVF-Ca^2+^ events with SMCs showing a consistent decrease in Ca^2+^ following whisker stimulation.

### Chemogenetic activation of brain fibroblast-Ca^2+^ signaling

Col1a2CreER-GqDREADDflox mice and littermate controls (GqDREADDfl/+ or Col1a2CreER/+) underwent ∼45 min of awake two-photon imaging to capture at least three distinct pial arteriole regions. Mice were then removed from the headmount holder and treadmill and intraperitoneally administered 100 μL Deschloroclozapine (DCZ; 30 μg/kg; Hello Bio, HB9126). Following DCZ administration, mice were re-secured to the headmount holder and treadmill, and repositioned under the two-photon microscope. The same vascular regions were subsequently reimaged for up to an additional hour. No additional isoflurane was administered during removal, DCZ injection, and remounting of mouse to avoid potential effects of anesthesia on vascular dynamics.

### Analysis of spatiotemporal dynamics of vasomotion

In movies of pial arteries from Col1a2CreER-GqDREADD and littermate controls, two ROIs of roughly equal distances (45-59 µm) were identified along continuous arterial segments lacking branchpoints. Only arterial segments that had continuous mCitrine-positive coverage, indicative of consistent PVF-GqDREADD expression along arteries in the Col1a2CreER-GqDREADD mice, were included in these analyses. To understand how well the oscillatory nature of ROIs A and B aligned pre and post DCZ, circular mean phase difference was calculated from Hilbert phase analysis using the hilbert() MATLAB function, and PLV between the two ROIs were calculated as described above. Mean coherence was calculated using magnitude-squared coherence via the mscohere() MATLAB function to assess the extent to which diameter oscillations were coupled between paired ROI regions within the range of vasomotor frequency.

### Intracisternal magna injections following DCZ administration

Col1a2CreER-GqDREADD and respective littermate controls were treated with DCZ (30µg/kg) 30 mins prior to intracisternal magna injections (ICM). About 15 mins prior to ICM injections, mice were anesthetized with a cocktail of ketamine (105mg/kg) and xylazine (15mg/kg) which better preserves CSF influx dynamics^24^. Prior to surgery, mice also received Carprofen (5.5mg/kg) and were mounted onto a stereotaxic frame. The back of the skull and scruff area was disinfected with betadine and 70% ethanol. A small incision was created to expose the posterior skull and muscle layers were carefully retracted using hemostats to reveal the cisterna magna located at the junction of the skull and spine. Four microliters of a mixture containing 1 mg/mL 45 kDa Ova-647 (Invitrogen; O34784) and 5 mg/mL 10 kDa FITC dextran (lysine fixable; Invitrogen; D1820) was injected using a 28 gauge Hamilton needle at a rate of 1 µL/min using a WPI microsyringe pump controller. After the 4 min injection, the needle was maintained in the cisterna magna for 1 min post injection to prevent backflow. The muscle and fascia were sutured, followed by the skin with 5-0 poly vicryl sutures. Mice were then removed from the stereotaxic frame and body temperature and anesthesia were maintained on a heating pad for the following 30 mins. Following ICM injections, mice were deeply anesthetized with pentobarbital and trans-cardial perfusions were performed with PBS followed by 4% paraformaldehyde. Brains were then dissected and prepared for immunohistochemistry.

### Immunohistochemistry following ICM injections

Brains were sectioned using a vibratome at 100 µm, collecting six equidistant sections per brain from bregma position 1.70 mm to -3.46mm. To label the vasculature, brain sections were stained with Lectin DyLight 594 (1:200; Vector Labs; DL-1177-1) in a solution of 2% TritonX-100, 5% BSA and 0.1% sodium azide in PBS for 48 hours at 4°C. Sections were washed 3 times for 5 minutes in PBS and mounted with Fluromount-G (ThermoFisher; 00-4958-02). Tile-scan immunofluorescent images at a magnification of 10x were captured of whole brain sections using a Zeiss 780 LSM confocal microscope.

### Analysis of CSF influx

All samples were blinded prior to analysis. Brain sections were analyzed in Fiji/ImageJ (v1.54p). Confocal z-stack images were converted to maximum intensity projections. For each section, the segmented line tool was used to manually outline and measure the total length of the cortical surface, as well as the regions of FITC-positive tracer signal. PAs were identified using the lectin staining and based on branching patterns shown in **Extended Data Fig. 9a**. Specifically, penetrating arterioles have few distinct off-shoots with a gradual decrease in diameter with subsequent branching while ascending venules generally have numerous off-shoots throughout the cortical column^25^. A few parenchymal vessels with overlapping or unclear branching patterns were classified as undefined and excluded from these analyses. Following the identification of PAs, they were measured using the segmented line tool. The length of detectable FITC tracer along the length of each PA was measured using the same approach. Cortical penetration depth of FITC tracer was assessed in PAs which were visible in entirety from the pial surface to the point of termination into a capillary network by measuring the maximum distance reached by the FITC tracer along the length of the vessel.

### Statistics

All statistical analyses were performed in GraphPad Prism (ver. 10). Respective statistical analyses are reported in each figure legend. Normality tests, generally Shapiro–Wilk tests, were performed on necessary datasets prior to statistical tests. For analyses reporting circular mean, Watson-Williams tests were performed, with Mardia-Watson-Wheeler tests used to compare overall circular distributions when low concentration (k) indicated the dispersion assumption of the Watson-Williams test was not met. SD is reported in all graphs unless specified otherwise in figure legends.

## Code availability

All custom code scripts are publicly available on https://github.com/StephanieKaras/PVF-VasomotionGitHub.

## ACKNOWLEDGMENTS

This work was supported by T32NS099042 to EH, 24POST1187589 to OB, T32AG052354 to CDN, grants to FD (R01HL136636, R01NS129022, RF1NS140137, Leducq Transatlantic Network of Excellence 22CVD01 BRENDA), grant to JAS (R01NS1311823), grants to AYS (R01AG081840, R01AG077731, R01NS097775, R01AG062738, Leducq Transatlantic Network of Excellence 23CVD03), and grants to SKB (R00AG080034, AARG-25-1488773, VCID-UMD-26-1529753). We would like to thank the Brain Clearance in CAA Leducq Foundation Network for their helpful insights and discussions during these studies. Special thanks to Dr. Susanne van Veluw for providing guidance during these studies.

## AUTHOR CONTRIBUTIONS

SKB, OB, JAS, and AYS conceptualized and designed experiments. Two-photon *in vivo* imaging was performed by SKB, SK, EH, and LMM. Development of *in vivo* analysis pipelines was done by SKB, SK, and OB. Analysis of *in vivo* datasets was performed by SKB, SK, EH, LMM, FL, and CDN. *Ex vivo* vessel preparations, experiments, and associated analyses were performed by DAJ and FD. Intracisterna magna injections and tissue collections were performed by SKB, SK, BP, LMM, MSI, and JAS. Immunostaining, confocal imaging, and respective analyses were performed by SK and EH. Statistics was performed by SKB, SK, and EH. Manuscript was written by SKB and SK with editing and contributions from all authors.

