## Supplemental Figures for "Perivascular fibroblasts locally regulate the vasomotor dynamics of pial arterioles"

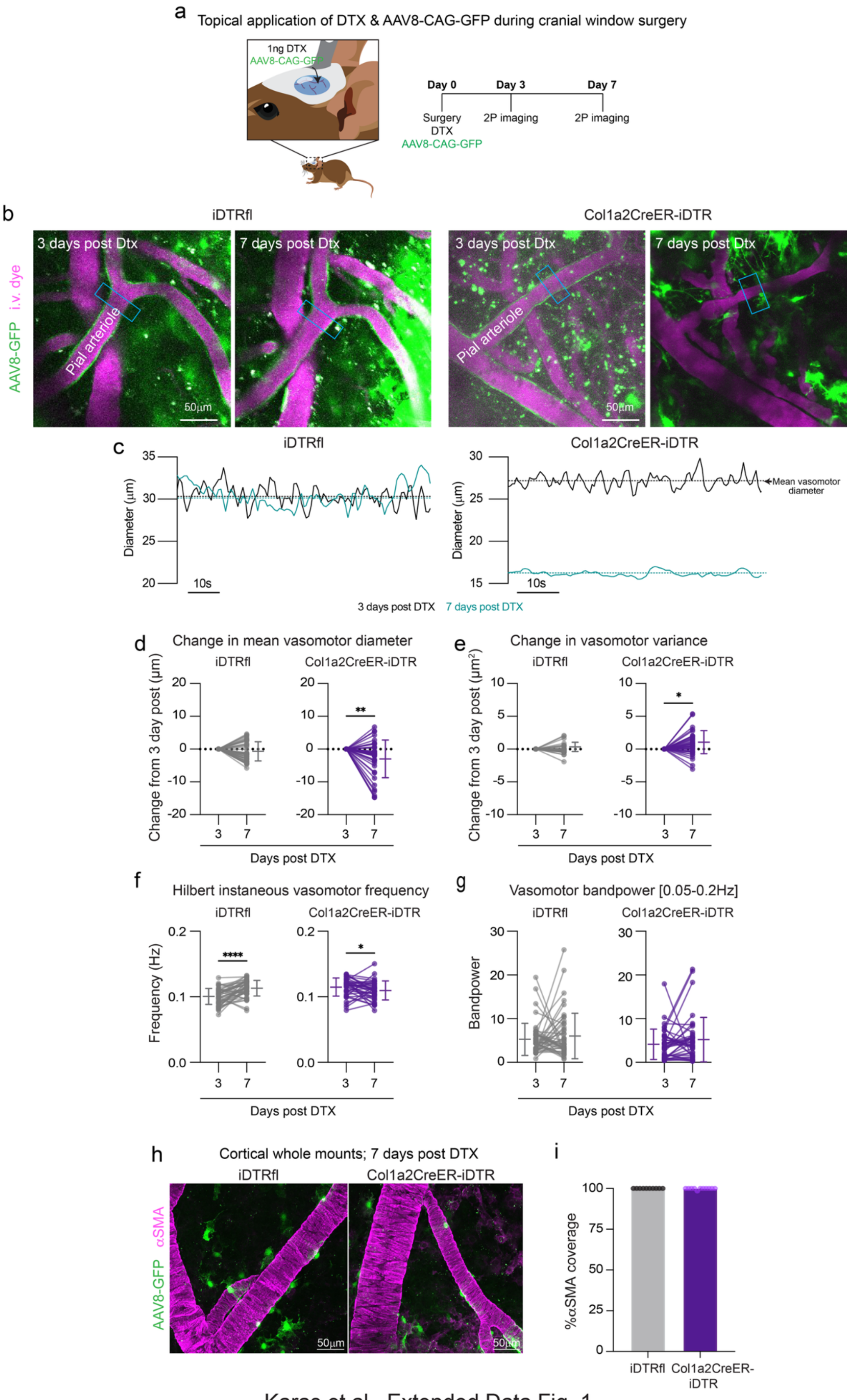


Karas et al., Extended Data Fig. 1

10

**Extended Data Figure 1. Brain fibroblast depletion disrupts vasodynamics for pial arterioles.**

**(a)** Graphical depiction of fibroblast labeling and depletion timeline following topical application of diphtheria toxin (DTX) and AAV8-CAG-GFP onto the surface of the brain in littermate controls (iDTRfl/+; iDTRfl) and Col1a2CreER-iDTR mice on Day 0 during skull-removed cranial window implantation. Animals then underwent imaging 3 and 7 days post DTX.

**(b)** Representative *in vivo* images of iDTRfl and Col1a2CreER-iDTR mice 3 and 7 days post DTX with an example of diameter region of interest (ROI) indicated in cyan box on pial arterioles. GFP puncta in LPMFs and PVFs, as indicated by the sheet of green cells around pial arterioles, appears 3 days post AAV8-CAG-GFP transduction. Seven days post DTX and AAV8-CAG-GFP application, we observe a reduction in PVFs surrounding pial arterioles and LPMFs in Col1a2CreER-iDTR mice.

**(c)** Graphs showing 1 min of resting diameter changes in the vascular territories from vessel regions shown above from iDTRfl and Col1a2CreER-iDTR with 3 (black) and 7 (green) days post DTX for each respective ROI. Dashed lines indicate mean vasomotor diameter.

**(d-g)** Graphs showing the **(d)** change in vasomotor diameter mean, **(e)** change in vasomotor variance, **(f)** Hilbert instantaneous frequency, and **(g)** vasomotor bandpower within 0.05-0.2 Hz 3 and 7 day post DTX for ROIs along pial arterioles in control (gray; iDTRfl) and Col1a2CreER-iDTR (purple) mice. Col1a2CreER-iDTR (n=3 mice) arteriolar ROIs n=38. Wilcoxon test 3 vs. 7 day DTX; change in vasomotor diameter mean **p=0.0047, mean instantaneous frequency *p=0.04. Paired t test 3 vs. 7 day DTX; change in vasomotor variance *p=0.0104. iDTRflox control mice (n=2 mice) arteriolar ROIs n=44. Paired t test 3 vs. 7 day DTX; mean instantaneous frequency ****p<0.0001.

**(h)** Representative confocal images of the cortical whole mounts of the cranial window from iDTRfl and Col1a2CreER-iDTR mice 7 days post DTX immunostained for αSMA and GFP.

**(i)** Graph showing quantification of percent (%) αSMA coverage on pial arterioles from cortical whole mounts of the cranial window from iDTR (n=3 mice) and Col1a2CreER-iDTR mice (n=3 mice) 7 days post DTX. Unpaired t-test iDTR vs. Col1a2CreER-iDTR; percent αSMA coverage p=0.3409. Each point is a separate image (3 per animal) taken at 20x within the cranial window area to show consistent αSMA coverage along pial arterioles with this depletion model.

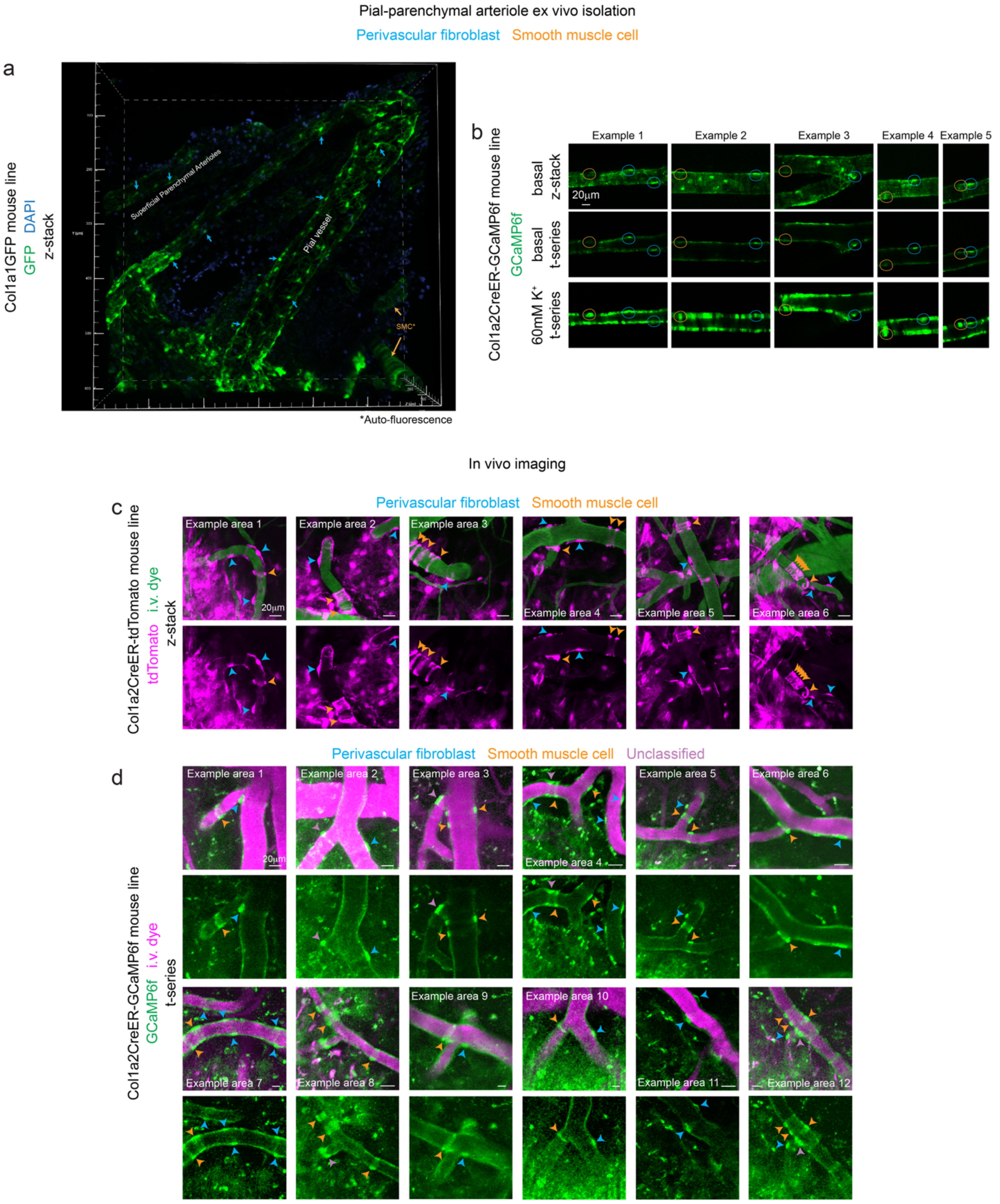


Karas et al., Extended Data Fig. 2

**Extended Data Figure 2. Morphological characterization of PVFs and SMCs *ex vivo* and *in vivo*.**

**(a)** Representative z-stack, max projected confocal image of isolated pial and parenchymal arteriolar networks isolated from Col1a1GFP mice. PVFs are indicated with blue arrows and exhibit flattened somata with thin, fibrillar processes along pial vessels and penetrating arterioles. Occasionally, autofluorescent SMCs are observed on isolated pial arterioles, characterized by their distinctive banding morphology (orange arrows). Nuclear labeling with DAPI shown in blue.

**(b)** Example *ex vivo* confocal images of a superficial parenchymal arterioles from Col1a2CreER-GCaMP6f mice with (top) z-stack, max projection taken under basal conditions, (middle) single-plane t-series under basal conditions, and (bottom) single-plane t-series immediately following 60mM $K^+$ exposure to the vasculature. GCaMP6f+ PVFs (blue arrows) are visible under basal conditions, with banding of recombined GCaMP6f-expressing SMCs (orange arrows) visible after 60mM $K^+$. Images shown here were purposefully chosen for a side-by-side comparison of PVFs and SMCs.

**(c)** Example *in vivo* z-stack, max projected images of tdTomato-positive (magenta) PVFs and SMCs on pial arterioles in Col1a2CreER-tdTomato mice. tdTomato-positive PVFs also exhibit flattened somata and thin, fibrillar processes (blue arrows). tdTomato-positive SMCs, indicated by orange arrows, wrap around the arterial wall and exhibit the characteristic banded morphology. Intravascular (i.v.) dye shown in green. Images shown here were purposefully chosen for a side-by-side comparison of PVFs and SMCs.

**(d)** Example *in vivo* single-plane t-series images of GCaMP6f-positive (green) PVFs and SMCs on pial arterioles in Col1a2CreER-GCaMP6f mice. GCaMP6f-positive SMCs (orange arrows) exhibited the banded morphology wrapping around the pial arterioles. In contrast, GCaMP6f-positive cells with flattened somata and lacking a banded morphology were classified as PVFs (blue arrows). A small number of cells exhibited overlapping morphological features and could not be classified, we therefore excluded these from our analyses (pink arrows). I.v. dye shown in magenta. Images shown here were purposefully chosen for a side-by-side comparison of PVFs and SMCs.

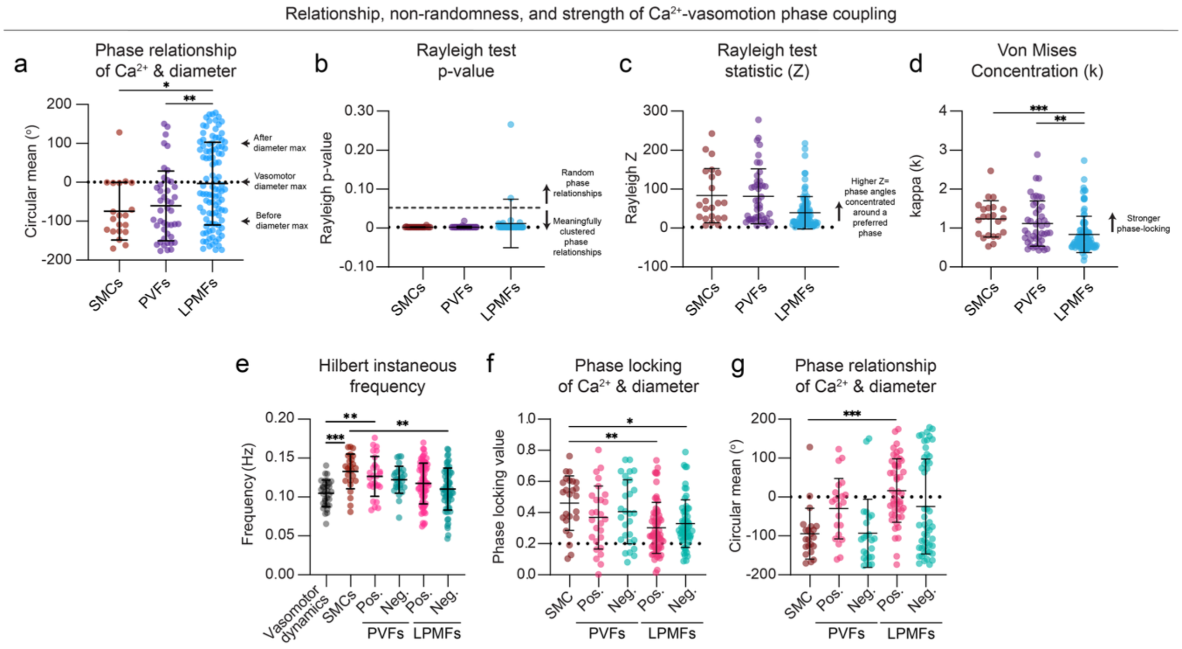


Karas et al., Extended Data Fig. 3

**Extended Data Figure 3. Comparison of $Ca^{2+}$-vasomotion phase metrics in SMCs, PVFs, and LPMFs.**

**(a-d)** Graphs reporting the relationship, non-randomness, and strength of $Ca^{2+}$-vasomotion phase coupling from Hilbert phase analyses showing **(a)** Preferred phase relationships of $Ca^{2+}$ with diameter (circular mean (°)), **(b)** Rayleigh test of uniformity p-value (Mean p-value: SMCs: p=0.0003, PVFs: p=0.0004, LPMFs: p=0.01) **(c)** Rayleigh test statistic (Z), and **(d)** Von Mises concentration (κ) comparing relationships between SMC- (red), PVF- (purple), and LPMF-$Ca^{2+}$ (blue) dynamics with diameter oscillations. SMC-$Ca^{2+}$ ROIs n=25, PVF-$Ca^{2+}$ ROIs n=65, LPMF-$Ca^{2+}$ ROIs n=155 in 3 Col1a2CreER-GCaMP6f mice. Circular means: Mardia-Watson-Wheeler test: SMCs vs. LPMFs: W=16.701, df=2, ***p=0.00071. Von Mises concentration: One-way ANOVA followed by Kruskal-Wallis test: SMCs vs. LPMFs: ***p=0.0003, PVFs vs. LPMFs: **p=0.0071. Graphs labeled to assist with interpretation of different analyses.

**(e)** Graph demonstrating Hilbert instantaneous frequency (Hz) of arteriolar vasodynamics (black) and SMC-$Ca^{2+}$ oscillations (red) as well as positive-dominant (pink) and negative-dominant (green) PVF-$Ca^{2+}$ and LPMF-$Ca^{2+}$ oscillations. Arteriolar ROIs = 32, SMC-$Ca^{2+}$ ROIs n=25, PVF-$Ca^{2+}$ ROIs n=68 (27 positive-dom., 28 negative-dom.), and LPMF-$Ca^{2+}$ ROIs n=155 (69 positive-dom., 61 negative-dom.) in 3 Col1a2CreER-GCaMP6f mice. Only significant comparisons versus SMCs and vasomotion (where applicable) are indicated in e-g. One-way ANOVA followed by Tukey's multiple comparison test: Vasomotor dynamics vs. SMC-$Ca^{2+}$: ***p=0.0003, Vasomotor dynamics vs. positive-dominant PVF-$Ca^{2+}$: **p=0.0083, SMC-$Ca^{2+}$ vs. negative dominant LPMF-$Ca^{2+}$: **p=0.0014.

**(f)** Graph comparing phase-locking of cellular-$Ca^{2+}$ and diameter oscillations for SMCs (red), positive-dominant (pink) and negative-dominant (green) PVFs and LPMFs. Same n values as reported in (e). One-way ANOVA followed by Kruskal-Wallis test: SMC-$Ca^{2+}$ vs. pos-dom. LPMF-$Ca^{2+}$: **p=0.0014, SMC-$Ca^{2+}$ vs. neg-dom. LPMF-$Ca^{2+}$: *p=0.0352.

**(g)** Graph of phase relationship (circular means (°)) between cellular-$Ca^{2+}$ and diameter oscillations for SMCs (red), positive-dominant (pink) and negative-dominant (green) PVFs and LPMFs. Cells with a PLV ≥0.2 (above dashed line in f). SMC-$Ca^{2+}$ ROIs: n=22, PVF-$Ca^{2+}$ ROIs: n=44 (21 positive-dom., 23 negative-dom.), LPMF-$Ca^{2+}$ ROIs: n=99 (48 positive-dom., 51 negative-dom.) in 3 Col1a2CreER-GCaMP6f mice. Mardia-Watson-Wheeler test: SMC-$Ca^{2+}$ vs. pos-dom. PVF-$Ca^{2+}$: W=10.828, df=2, *p=0.027, SMC-$Ca^{2+}$ vs. pos-dom. LPMF-$Ca^{2+}$: W=26.186, df=2, ****p=0.000016.

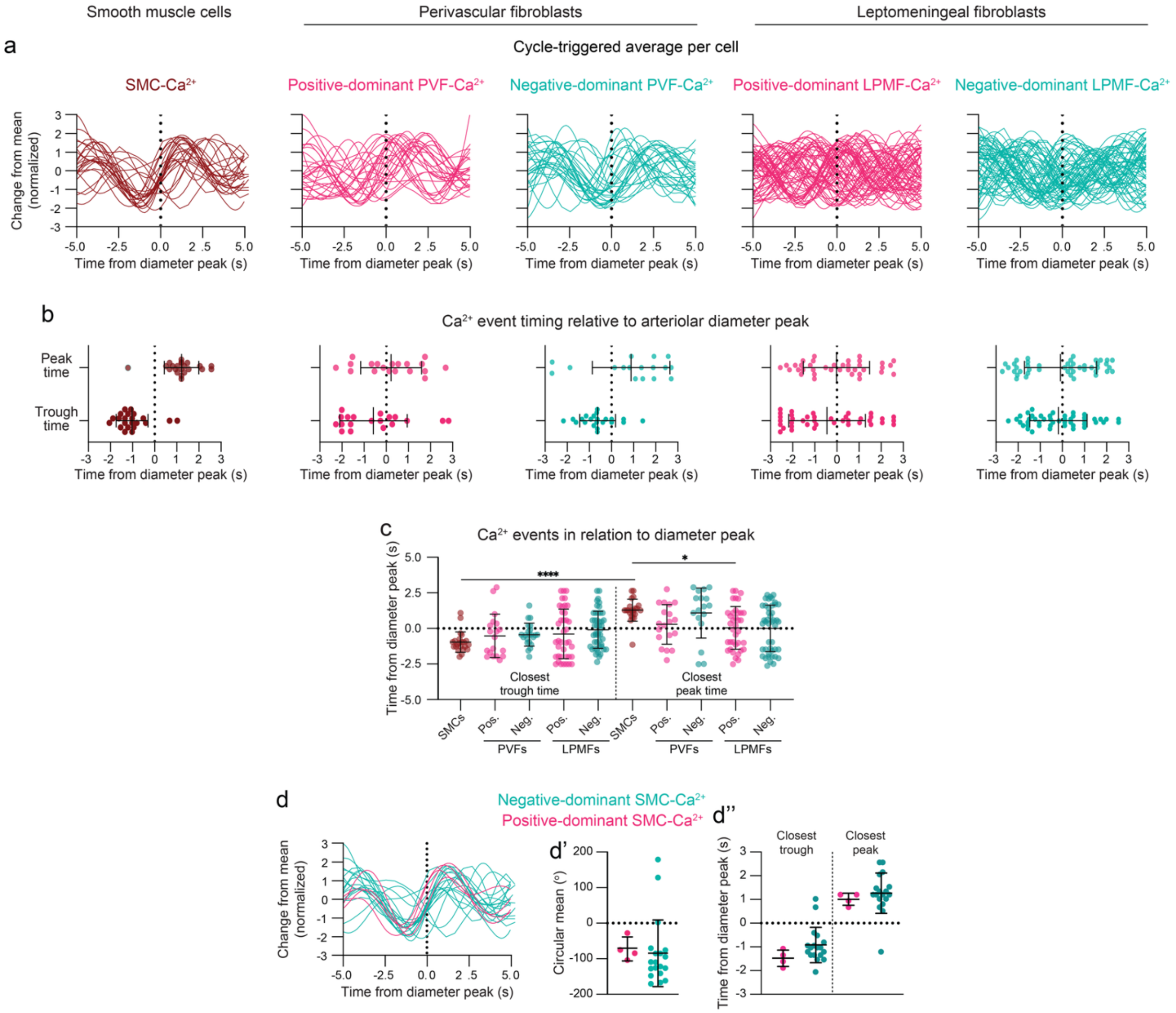


Karas et al., Extended Data Fig. 4

**Extended Data Figure 4. $Ca^{2+}$ dynamics of individual cells show unique relationships within individual vasomotor cycles from cycle-triggered analyses.**

**(a)** Cycle-triggered average plots based on average vasomotor cycles with peak dilation represented at 0 seconds. Individual SMC-$Ca^{2+}$ (red), along with positive-dominant-$Ca^{2+}$ (pink) and negative-dominant $Ca^{2+}$ (green) PVFs and LPFMs. Each cycle represents a 10 s vasomotor oscillation (0.1Hz). All cells have a phase lock value ≥0.2. SMC-$Ca^{2+}$ ROIs: n=22, PVF-$Ca^{2+}$ ROIs: n=44 (21 positive-dom., 23 negative-dom.), LPMF-$Ca^{2+}$ ROIs: n=99 (48 positive-dom., 51 negative-dom.) in 3 Col1a2CreER-GCaMP6f mice.

**(b)** Graphs of the closest trough and peak times for each cell time in relation to diameter peak centered around 0 s.

**(c)** Graph comparing cellular $Ca^{2+}$ trough and peak events between each fibroblast correlation group with diameter peak centered around time zero. One-way ANOVA followed by Kruskal-Wallis test: Closest $Ca^{2+}$ trough vs. peak - SMCs: ****p<0.0001, PVFs: *p=0.0135. SMCs vs. positive-dominant LPMFs - Closest $Ca^{2+}$ peak: *p=0.0352.

**(d)** Graphs of **(d)** cycle-triggered average, **(d')** circular means (°), and **(d'')** closest $Ca^{2+}$ trough and peak, comparing negative-dominant and positive-dominant SMC-$Ca^{2+}$ relationships with vasomotor dynamics showing slight shift in phase relationships and $Ca^{2+}$ dynamics closer to peak dilation for SMCs classified as positive dominant (positive r value). This demonstrates that that correlation sign reflects differences in temporal positioning within the vasomotor cycle rather than fundamentally distinct calcium behaviors.

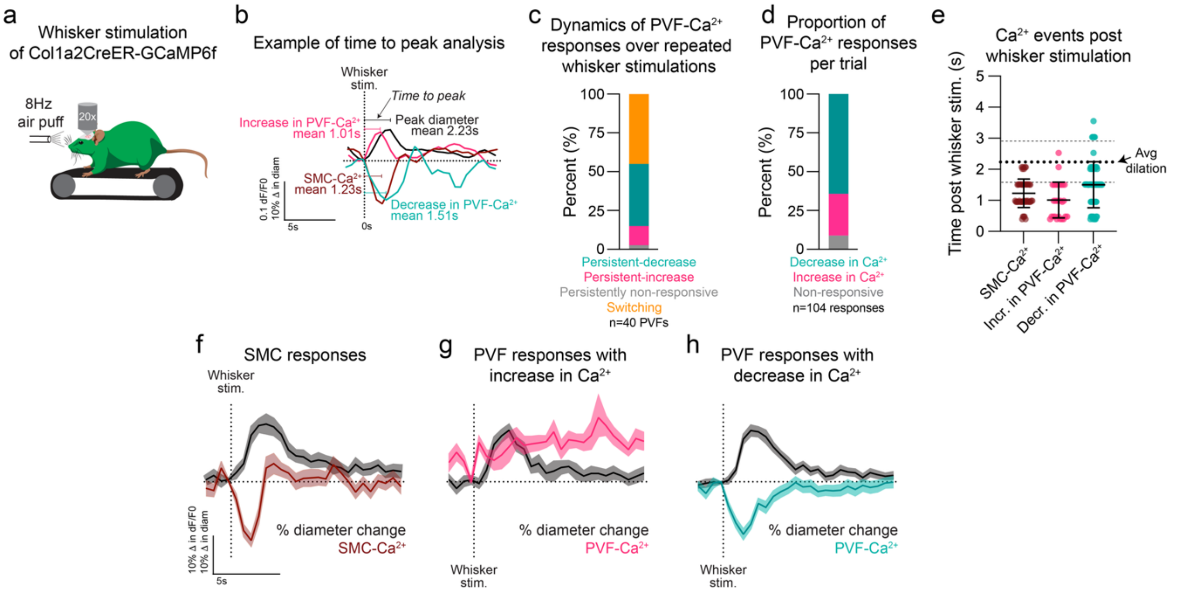


Karas et al., Extended Data Fig. 5

**Extended Data Figure 5. PVFs display similar $Ca^{2+}$ relationships to SMCs during neuronal-evoked vasodilation of pial arterioles.**

**(a)** Graphical depiction of awake in vivo imaging setup of Col1a2CreER-GCaMP6f mice head-fixed on freely-moving treadmill along during 8Hz air puff whisker stimulation.

**(b)** Graphical depiction of time to peak analysis post whisker stimulation on SMC-$Ca^{2+}$, PVF-$Ca^{2+}$, and diameter responses. Examples of an increase in PVF-$Ca^{2+}$ are shown pink while a decrease in PVF-$Ca^{2+}$ post whisker stimulation is shown in green. SMCs (red) were identified by their banded morphology and consistently displayed a decrease in $Ca^{2+}$ following whisker stimulation. Vascular responses due to whisker stimulation are shown in black. All mean time to peak responses are indicated for each perivascular cell subtype and vessel dilation.

**(c)** Graph showing the $Ca^{2+}$ dynamics from single PVF ROIs over repeated whisker stimulations. PVFs with increase in $Ca^{2+}$ post whisker stimulation over the multiple trials were defined as having a persistent-increase (pink) with PVFs displaying a persistent-decrease in $Ca^{2+}$ are shown in green. Switching PVFs (orange) were defined as PVFs that exhibited both positive and negative $Ca^{2+}$ responses across multiple whisker stimulation trials. Cells with no $Ca^{2+}$ response post whisker stimulation over the repeated trials were classified as persistently non-responsive (gray). A total of 40 PVFs were analyzed, n=2 mice, 3-5 areas analyzed per animal.

**(d)** Graph showing the proportion of PVF-$Ca^{2+}$ responses per trial based on an increase in $Ca^{2+}$ (pink), decrease in $Ca^{2+}$ (green), or non-responsive (gray). A total of 104 PVF-$Ca^{2+}$ responses were analyzed.

**(e)** Graph showing the time to first $Ca^{2+}$ event post whisker stimulation for SMC-$Ca^{2+}$ responses (n=48), positive-dominant PVF events (n=28), and negative-dominant PVF events (n=67).

Average dilation time indicated with the dashed line at 2.23 s with standard deviation of +/-0.66 seconds with the thinner dashed lines.

**(f-h)** Graphs showing the mean $Ca^{2+}$ responses from **(f)** SMCs, **(g)** positive-dominant PVFs, **(h)** negative-dominant PVFs post whisker stimulation alongside their respective mean dilations of their vascular territories. SEM indicated for mean $Ca^{2+}$ and diameter.

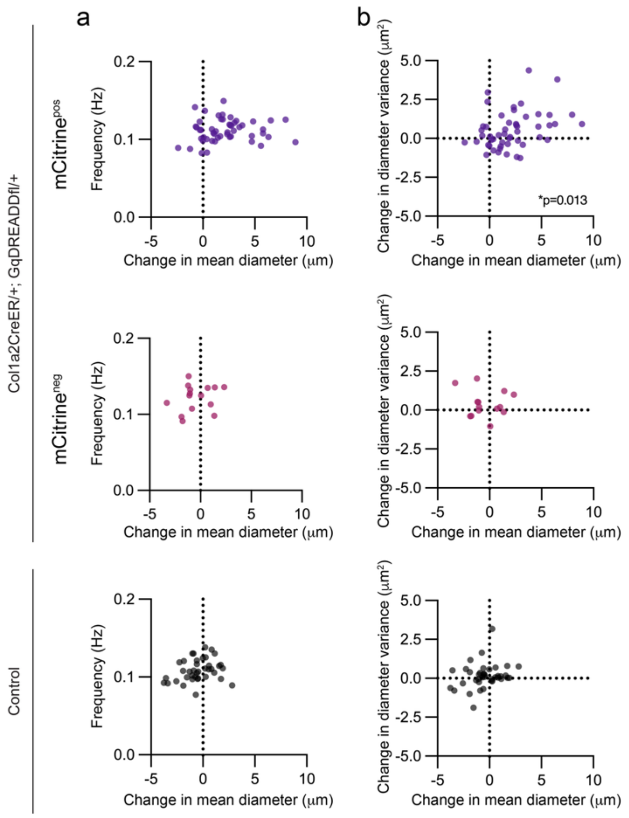


Karas et al., Extended Data Fig. 6

**Extended Data Figure 6. Activation of perivascular fibroblast-Gq signaling enhances arteriolar vasomotor dynamics.**

**(a & b)** Correlation plots of change in mean diameter vs **(a)** frequency (Hilbert instantaneous frequency) and **(b)** change in diameter variance in control mice (GqDREADDfl/+ or Col1a2CreER/+; black) or mCitrine-positive (purple; mCitrine$^{pos}$) and mCitrine-negative (pink; mCitrine$^{neg}$) vessel segments in Col1a2CreER-GqDREADD mice post DCZ (40-60min post DCZ). Potential relationships analyzed using Pearson correlation tests with significant correlations reported on respective graphs.

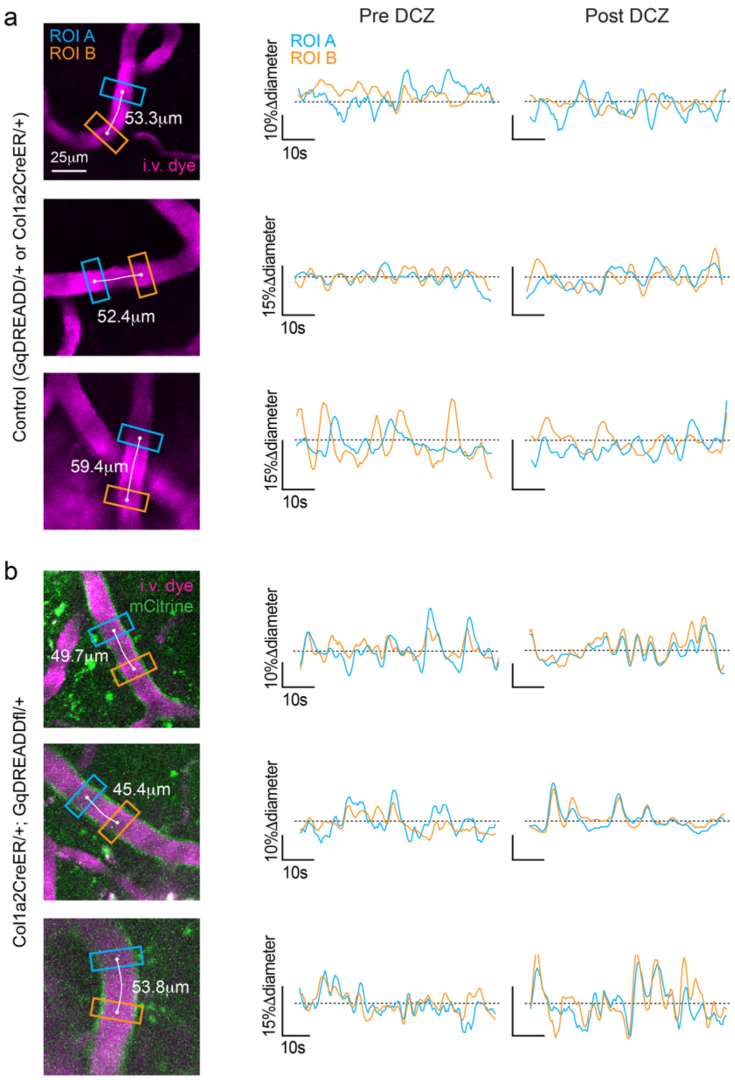


Karas et al., Extended Data Fig. 7

**Extended Data Figure 7. Examples of improved spatiotemporal vasodynamics of pial arterioles in Col1a2CreER-GqDREADD mice post DCZ.**

**(a & b)** Three representative in vivo images from **(a)** control (GqDREADDfl/+ or Col1a2CreER/+) and **(b)** Col1a2CreER-GqDREADD with their respective regions of interest (ROI) A (blue) and B (orange) with vessel distance around 50μm. Each image accompanied by respective percent diameter changes for each ROI overlaid pre and post DCZ during 1 min of rest to demonstrate increased synchrony following activation of PVF-Gq signaling.

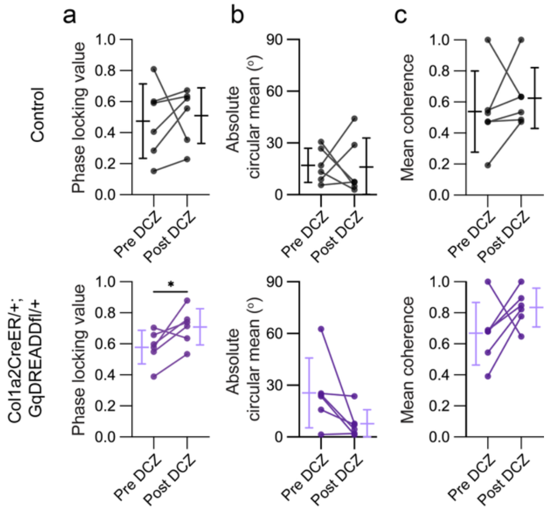


Karas et al., Extended Data Fig. 8

**Extended Data Figure 8. Activation of perivascular fibroblast-Gq signaling improves the spatiotemporal vasodynamics of pial arterioles in Col1a2CreER-GqDREADD mice.**

**(a-c)** Graphs showing the synchrony of ROI A vs ROI B via **(a)** phase-locking value, **(b)** Absolute circular mean ($^{o}$), and **(c)** mean coherence in control (GqDREADDfl/+ or Col1a2CreER/+; black) and Col1a2CreER-GqDREADD (purple) mice. Control analysis (n=6 pial arterioles; 3 mice) - pre vs post DCZ, Paired t-test: phase-locking value p=0.74, Mean coherence p=0.46. Col1a2CreER-GqDREADD analysis (n=6 pial arterioles; 3 mice) - pre vs post DCZ, Paired t-test: phase-locking value *p=0.045, Mean coherence p=0.17. Mardia-Watson-Wheeler test: absolute circular mean ($^{O}$) W=4.88, df=2, p=0.087.

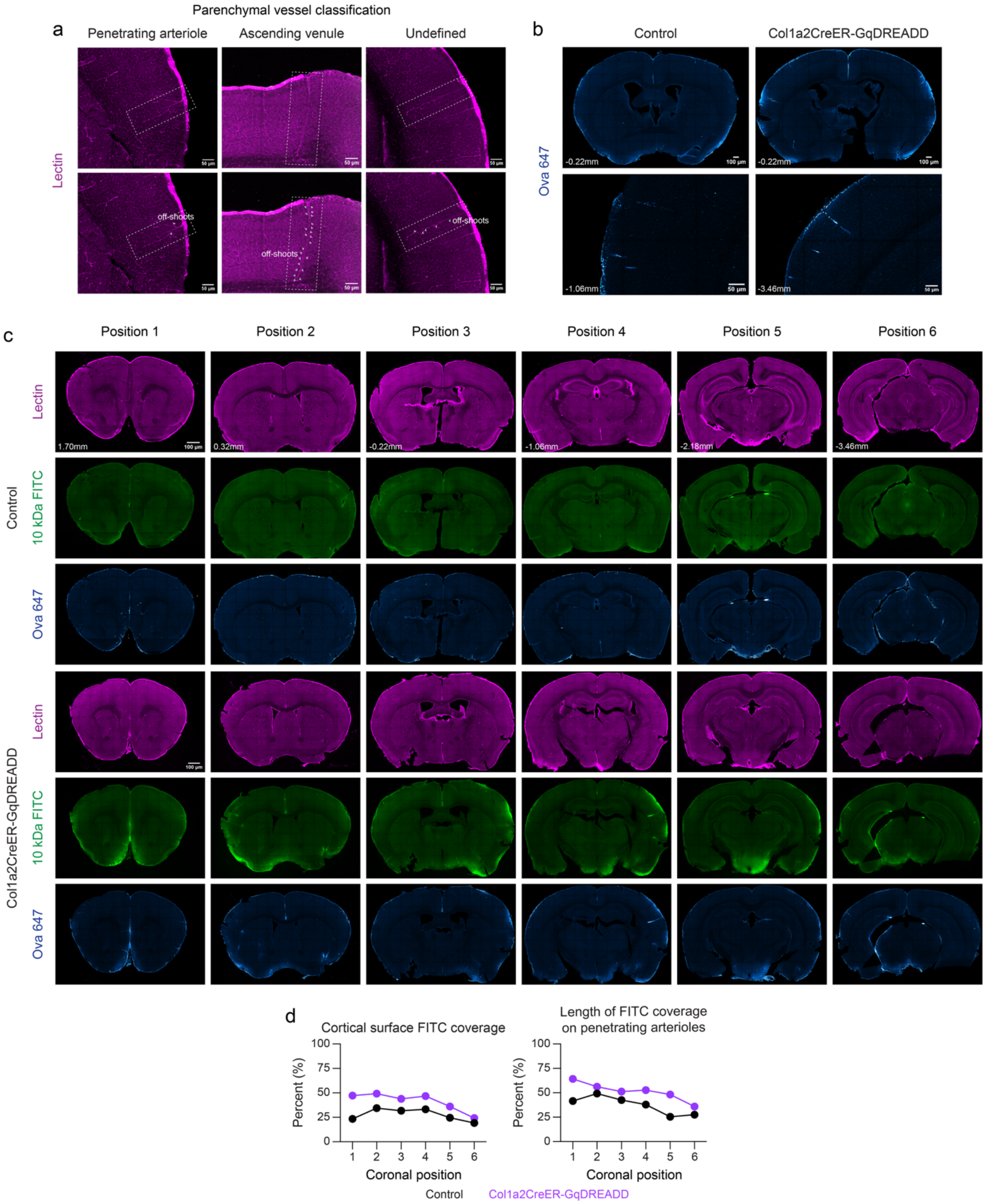


Karas et al., Extended Data Fig. 9

184 **Extended Data Figure 9. Characterization of CSF influx following chemogenetic activation**
185 **of fibroblast-Gq signaling.**

**(a)** Representative confocal images of brain sections stained for Lectin (magenta) illustrating the morphological classification of parenchymal arterioles and venules. Penetrating arterioles (left) exhibit few distinct off-shoots (white arrows) that decrease in diameter gradually with each subsequent branching event. This distinguishes them from ascending venules (middle), which exhibit multiple off-shoot vessels. A few vessels within these experiments demonstrating overlapping or unclear morphological features were classified as undefined (right) and excluded from analyses.

**(b)** Representative cortical images corresponding to the regions shown in Fig. 3 of 10kDa-FITC dextran showing similar distribution of Ova-647 (blue) in control (GqDREADDfl/+ or Col1a2CreER/+) and Col1a2CreER-GqDREADD mice following intracisternal magna injections. Corresponding positions relative to bregma noted.

**(c)** Representative coronal sections of the six equidistant positions analyzed per mouse, as defined by bregma coordinates, ordered from most rostral (position 1; 1.70mm) to most caudal (position 6; -3.46mm). Sections are shown for control and Col1a2CreER-GqDREADD mice with lectin labeling (magenta), 10 kDa FTIC tracer (green), and Ova-647 (cyan).

**(d)** Graphs of FITC tracer coverage on the cortical surface (left) and along the length of penetrating arterioles (right) across the six rostral to caudal coronal positions analyzed per mouse (n=6 control, n=5 Col1a2CreER-GqDREADD). Points represent group means at each position. Two-way ANOVA for FITC tracer coverage on the cortical surface: significant main effect of cortical position *p=0.0372 and genotype **p=0.003. FITC tracer on length of penetrating arterioles: significant main effect of cortical position *p=0.0398 and genotype **p=0.005.
